# Bayesian Causal Inference Accounts for Multisensory Filling-In at the Blind Spot

**DOI:** 10.1101/2024.11.15.623713

**Authors:** Ailene Y. C. Chan, Noelle R. B. Stiles, Carmel A. Levitan, Daw-An Wu, Armand R. Tanguay, Shinsuke Shimojo

**Affiliations:** California Institute of Technology, Division of Biology and Biological Engineering; Rutgers University, Departments of Neurology, Ophthalmology and Visual Science, Biomedical Engineering; Center for Advanced Human Brain Imaging Research in the Brain Health Institute; Occidental College, Cognitive Science; University of Southern California, Departments of Electrical Engineering, Chemical Engineering and Materials Science, Biomedical Engineering, Ophthalmology, and Physics and Astronomy; Neuroscience Graduate Program

**Keywords:** Vision, Blindspot, Audition, Multisensory, Filling-in, Illusion

## Abstract

We asked three questions about multisensory perception across the physiological blind spot: (1) Does audiovisual integration persist without bottom-up visual input? (2) Does the brain adjust its sensory uncertainties and priors accordingly? (3) Are the underlying causal-inference computations preserved?

Participants judged flashes and beeps in an audiovisual illusion presented across the blind spot or a matched control location. Responses were fit with a Bayesian Causal Inference (BCI) model, estimating sensory noise, numerosity priors, and causal-inference priors under multiple decision strategies evaluated using BIC.

Illusions were robust at both locations, indicating preserved integration. Model fits showed higher visual uncertainty and broader prior expectations at the blind spot, while auditory precision and the causal prior remained stable. Thus, the computational architecture of causal inference is maintained, but its parameters flexibly adapt to local sensory reliability.

These findings demonstrate that perceptual inference remains intact even in regions without retinal input, achieved by adjusting internal uncertainty rather than altering core multisensory computations.

## 1 Introduction

Perception feels continuous even though sensory inputs are incomplete and unevenly reliable. A major view in perceptual science is that the brain maintains this stability through Bayesian inference, using learned Bayesian priors—internal models of the world—to interpret incoming signals (Knill and Richards, 1996). Some of the clearest evidence for such priors comes from the physiological blind spot, a retinal region with no photoreceptors, where the brain seamlessly “fills in” missing visual information (Ramachandran, 1992). In the complete absence of feedforward input, top-down predictions dominate, and the content that is filled in reflects the structure of the brain’s learned internal model rather than external stimulation. Classic accounts emphasize mechanisms such as visual interpolation and predictive coding and thus imply that blind-spot filling-in is supported by strong, highly structured visual priors (Devinck and Knoblauch, 2019; Raman and Sarkar, 2016; Weil and Rees, 2011). Critically, these accounts remain almost exclusively unimodal: they explain how the visual system restores missing visual content, but they do not address how the brain resolves multisensory events that traverse a region with no bottom-up visual signal.

This makes the blind spot an exceptionally powerful test case for multisensory perception: a region with intact spatial and temporal context but no visual input. If perception is guided by stable Bayesian priors about how audiovisual events unfold, do these priors extend to a location where the visual likelihood collapses? One possibility is that multisensory effects should be stronger across the blind spot, reflecting greater reliance on crossmodal cues and a tendency to “fill in” missing information. Another possibility is that filling-in aims to maintain perceptual continuity, in which case multisensory effects should remain unchanged, matching those in fully visible regions.

To answer this, we used the Audiovisual (AV) Rabbit Illusion (Stiles et al., 2018), a spatial-temporal variant of the Double Flash Illusion (Shams et al., 2002). In the Double Flash Illusion, a single flash paired with two beeps leads observers to perceive two flashes, demonstrating that auditory information can alter visual temporal perception. The AV Rabbit Illusion extends this phenomenon: when two flashes are paired with three beeps, participants typically perceive an illusory flash between the two real flashes. Conversely, when three flashes are paired with only two beeps (the “invisible” variant), the middle flash is perceptually suppressed. This illusion is ideal for probing inference because its percept depends on the brain’s assumed causal structure linking auditory and visual events. We used a rigorous quantitative tool (Bayesian Causal Inference; BCI) to investigate the underlying computations of multisensory perception (Körding et al., 2007; Shams, 2012; Aller and Noppeney, 2019; Shams and Beierholm, 2022). If the brain applies the same Bayesian priors across space, illusion strength and model parameters should be identical across the blind spot and a matched control location. If, however, the brain adapts these priors when vision provides no input, multisensory perception should be preserved phenomenologically but supported by different computational signatures—for example, broader visual likelihoods or altered sensory priors.

Participants viewed audiovisual sequences either crossing the blind spot or a visible control region. Although illusion strength was remarkably similar across locations, BCI modeling revealed systematic differences in the underlying computations. Across the blind spot, observers exhibited broader visual likelihoods and flatter sensory priors, indicating that the brain maintains perceptual continuity not by applying the same priors used for visible regions but by flexibly adjusting them when bottom-up visual evidence is absent. Notably, the inferred causal structure—the tendency to bind auditory and visual events—did not differ across locations. Instead, the broader visual likelihoods and flatter priors at the blind spot increase the relative influence of auditory signals on the final percept. These findings suggest that blind-spot filling-in extends beyond vision: the brain engages a form of multisensory filling-in, preserving perceptual continuity through location-specific Bayesian inference even when one modality provides no direct input.

## 2 Methods

### 2.1 Human participants

Twenty participants (10 females; mean age = 26.7 years, range = 19–40 years) took part in the study. The sample size was determined based on prior power calculations using G*Power. All participants had normal or corrected-to-normal vision (20/20) and normal hearing, were naïve to the purpose of the study, and provided written informed consent before participation. Participants were compensated at $25/hour. All procedures were approved by the institutional review board at the California Institute of Technology (IR19-0926).

### 2.2 Stimuli and conditions

#### 2.2.1 AV Rabbit Task

Visual stimuli were white bars (0.28^°^ × 1.2^°^ visual angle) presented on a black background (100% contrast) for one frame (16.7 ms at 60 Hz). Auditory stimuli were pure tones (800 Hz, 7 ms). Six audiovisual conditions were tested using the naming convention FXBY, where X is the number of flashes and Y the number of beeps: (i) Unimodal (visual only): F2B0, F3B0, (ii) Bimodal congruent: F2B2, F3B3, (iii) Bimodal incongruent: F2B3 (Illusory AV Rabbit) and F3B2 (Invisible AV Rabbit) Stimulus timing and spatial configurations are illustrated in Fig. 1A–B. Stimuli were presented either across the blind spot or a corresponding control location matched in retinal coordinates. On each trial, flashes followed either a horizontal or vertical trajectory. For example, in a F3B0 trial, flashes could appear sequentially left–center–right or top–center–bottom; in a F2B0 trial, only the border positions (e.g., left–right) were shown.

**Figure 1:**
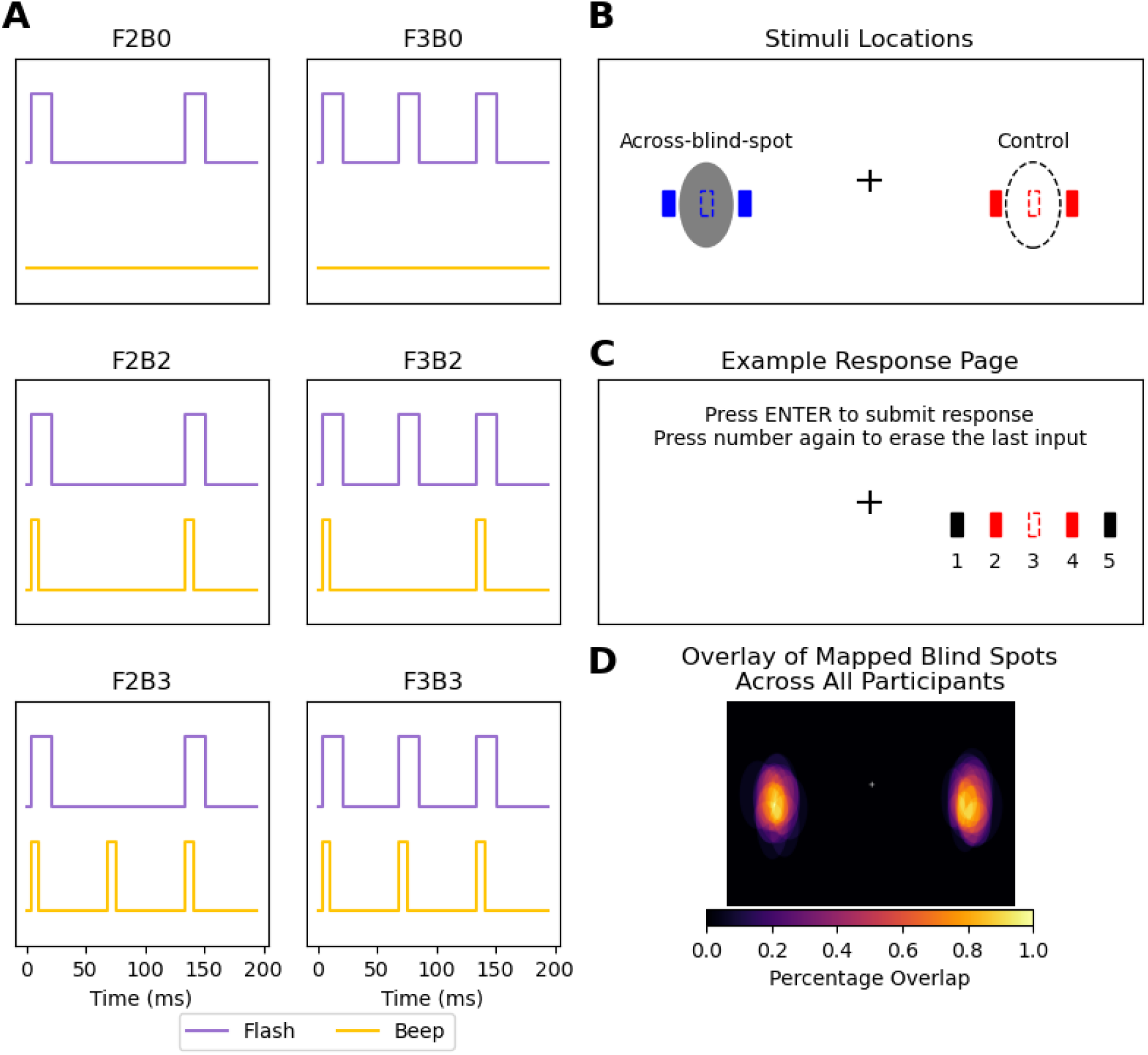
Experimental Design. **(A)** Schematic of the six experimental conditions, named using an FXBY convention (*X* = number of flashes, *Y* = number of beeps). When paired, visual flashes (16.7 ms) and auditory beeps (7 ms) had simultaneous onsets. Key conditions include the **Illusory AV Rabbit (F2B3)**, where an illusory flash is perceived with the second beep, and the **Invisible AV Rabbit (F3B2)**, where the middle flash is perceptually suppressed by the missing second beep. **(B)** Spatial configuration of stimuli shown for the left eye. Flashes were presented at the center of the blind spot (grey area) or a corresponding control spot (hollow area), as well as 0.5 visual degrees from the cardinal borders of these spots. In *F2*BY conditions, the two flashes appeared at the solid border points. In *F3*BY conditions, the first and third flashes appeared at the borders, while the second flash was presented at the center of the blind spot or control spot. For illustration, flashes traveling across the blind spot and control spot are colored blue and red, respectively; all flashes in the experiment were white. **(C)** The response interface for AV Rabbit Task. Participants reported the number of flashes they perceived by selecting from five equidistant options. Multiple selections were permitted. The colors are for illustrative purposes only. The response interface for the *Additional Flash Localization Task* is shown in Supplementary Video 1. **(D)** Overlay of the mapped blind spots across participants (N = 20). Blind-spot locations and sizes were consistent across individuals (see Fig. S1).

#### 2.2.2 Blind Spot Mapping

The blind spot of each eye was mapped individually using a flickering circular target (0.5°) alternating between black and white at 7 Hz on a black background. Participants monocularly fixated while moving the target to determine the disappearance and reappearance points along the cardinal directions (left, right, top, bottom). The same locations in the fellow eye’s visual field served as control spots, ensuring that stimuli at both positions occupied equivalent spatial coordinates within the visual field.

#### 2.2.3 Rationale for Control Spots

By using the fellow eye’s blind spot coordinates as control locations, the experiment compared perceptual performance for identical visual-field positions — once when all flashes were visible (control) and once when one flash fell within the blind spot (partial visual input). For instance, when the left eye was tested, the control spot corresponded to the right eye’s blind spot location but was fully visible to the left eye.

### 2.3 Experimental design

#### 2.3.1 Blind spot mapping protocol

Each eye was mapped in three short trials. Participants reported when the flickering target disappeared and reappeared while moving it horizontally or vertically. In trial 1, horizontal borders (left/right) were marked; in trial 2, vertical borders (top/bottom); and in trial 3, horizontal borders were rechecked. Fixation was monitored via an EyeLink 1000 eye tracker. After mapping, the estimated blind spot outline was displayed for participant verification. If borders appeared fuzzy, mapping was accepted; if they appeared sharp, the procedure was repeated.

#### 2.3.2 AV Rabbit Task

Participants completed two blocks: one reporting flash perception and one reporting beep perception. Each trial began with 500 ms of stable fixation (verified by eye tracking), followed by the audiovisual sequence and a 500 ms delay before response. In the flash block, participants selected the perceived number and positions of flashes using five equidistant options on screen (Fig. 1C). In the beep block, they reported the number of beeps. The relationship between flashes and beeps was never explained.

The design followed a 2 × 3 × 4 × 2 factorial structure: (i) flashes (2, 3), (ii) beeps (0, 2, 3), (iii) trajectory direction (horizontal/vertical × 2 directions), (iv) stimulus location (across-blind-spot, control), yielding 48 conditions. Each condition was repeated five times per block (240 trials per block), presented in random order.

#### 2.3.3 Additional Flash Localization Task

A subset of 10 participants returned for additional F2B3 (Illusory Rabbit) localization trials. Each eye was tested separately with 8 trials per eye (4 blind spot, 4 control). Participants, maintaining central fixation, used a crosshair cursor to click perceived flash positions on the screen. The position of the first flash was shown as a reference (see Supplementary Video 1).

### 2.4 Experimental setup

Stimuli were presented using Psychtoolbox 3.0.19 in MATLAB 2024a. Auditory tones ( 62 dB SPL) were delivered via stereo speakers (Bose Companion 2 Series III) placed beside the monitor. Visual stimuli appeared on a 27-inch LCD display (Dell UltraSharp U2720Q; 60 Hz). Eye position was recorded monocularly at 500 Hz with an EyeLink 1000 (SR Research, Canada) in remote mode. Participants sat 50 cm from the display with their head stabilized on a chin rest and the non-tested eye occluded. The experiment took place in a darkened room. All experimental scripts are publicly available under a CC-BY license (Chan, 2025).

### 2.5 Data analysis

#### 2.5.1 Accuracy

Accuracy was defined as the proportion of trials in which participants correctly reported the true number of flashes and beeps presented (Fig. 2A). To test whether the number of beeps influenced flash perception (i.e., the occurrence of AV Rabbit Illusions), we analyzed accuracy across beep conditions (B0, B2, B3) separately for two- and three-flash trials. For each flash condition and location, accuracy differences across beep levels were assessed using Friedman tests with Bonferroni-corrected Wilcoxon signed-rank post hoc comparisons. Within each condition, accuracy between the blind-spot and control locations was compared using Wilcoxon signed-rank tests (Bonferroni-corrected for multiple comparisons).

**Figure 2:**
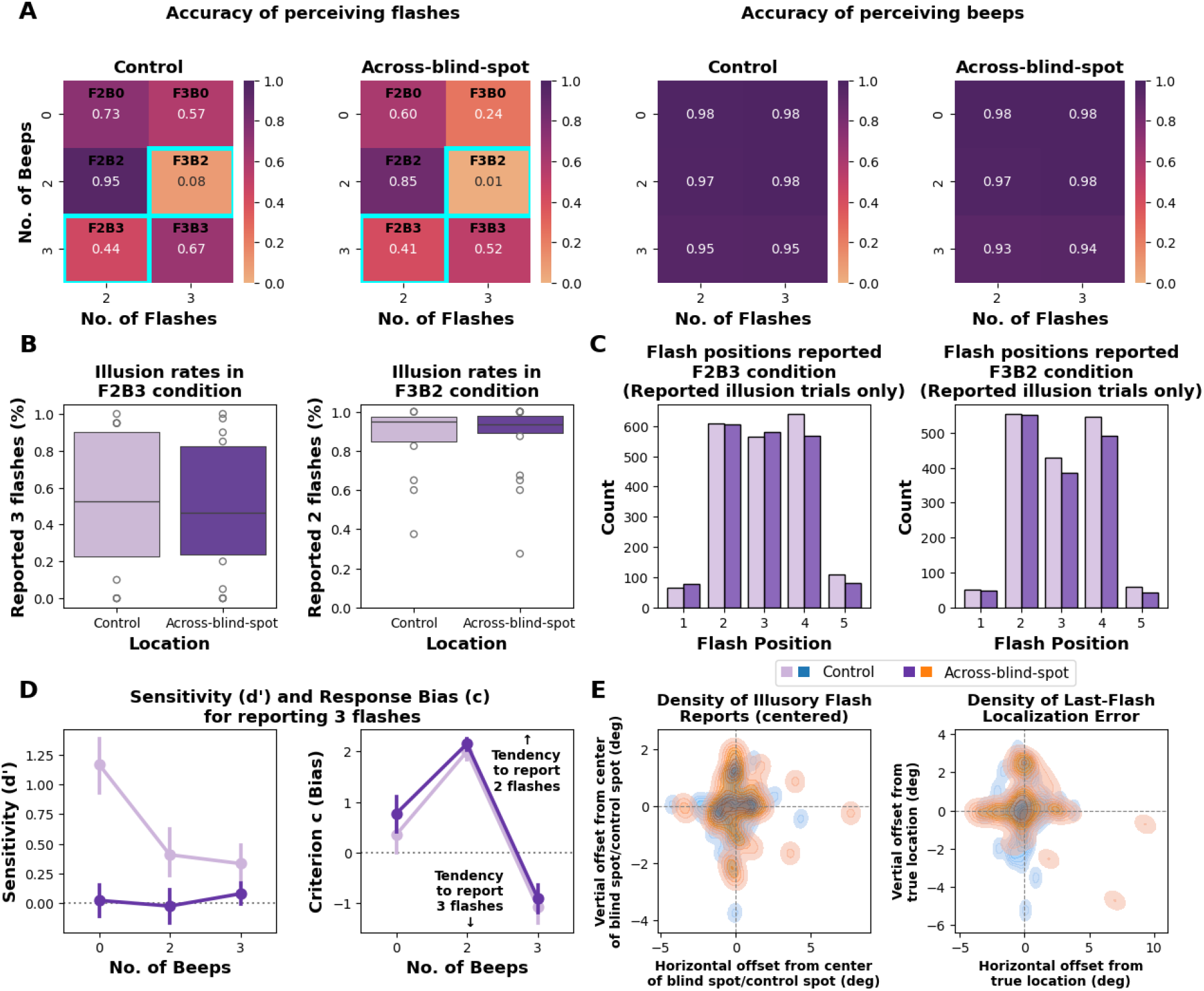
Effect of beeps on flash perception. **(A)** Accuracy for reporting flashes (left) and beeps (right) across control and across-blind-spot stimulus locations. Flash accuracy decreased markedly in conditions where the number of flashes and beeps mismatched (F2B3 and F3B2). Participants also showed reduced accuracy for three-flash stimuli presented across the blind spot in the unimodal (F3B0) condition. Beep-report accuracy remained near ceiling across all conditions. **(B)** Illusion rates in the Illusory Audiovisual Rabbit (F2B3) and Invisible Audiovisual Rabbit (F3B2) conditions were comparable between stimulus locations. **(C)** Reported flash positions in the illusion conditions. In F2B3, the extra beep induced an illusory flash between the two real flashes; in F3B2, the missing beep suppressed perception of the middle flash between the first and third flashes. **(D)** Signal detection results for reporting three flashes. Sensitivity (*d*^*′*^) was higher at control than across-blind-spot locations. Response criterion (*c*) was similar between locations, with two beeps biasing reports toward two flashes and three beeps toward three flashes. **(E)** Two-dimensional kernel density maps showing spatial offsets (in degrees of visual angle) of perceived illusory flashes (left) and last-flash reports (right). Offsets are computed relative to the center of the blind-spot/control region for illusory flashes, and relative to the true physical location of the last flash for last-flash localization errors. Horizontal and vertical offsets represent signed deviations along each axis.

#### 2.5.2 Illusion Frequency

To quantify the strength of the two AV rabbit illusions, we computed for each participant and location the proportion of illusion responses (Fig. 2B): *P* (respond 3| *F*2*B*3) for the Illusory AV Rabbit and *P* (respond 2| *F*3*B*2) for the Invisible AV Rabbit. These proportions were compared between control and across-blind-spot locations using Wilcoxon signed-rank tests (Bonferroni-corrected for multiple comparisons).

#### 2.5.3 Perceived Flash Position

For each AV rabbit illusion, we analyzed the distribution of reported flash positions (1–5) in illusion trials only (Fig. 2C). Biases in perceived position were first evaluated with chi-square goodness-of-fit tests against a uniform distribution, and the distributions between control and across-blind-spot locations were compared using chi-square tests of independence.

#### 2.5.4 Signal Detection Theory

To quantify perceptual sensitivity and decision criterion under each condition, we applied signal detection theory (SDT) separately for the Illusory AV Rabbit (3 beeps; B3), Invisible AV Rabbit (2 beeps; B2), and no-beep baseline (0 beeps; B0) contexts (Fig. 2D).

In each beep context, trials with three flashes (F3) served as *signal* and trials with two flashes (F2) as *noise*. For each participant and location (across-blind-spot, control), we computed the hit rate *H* = *P* (respond “3”| *F*3, *B*) and false alarm rate *FA* = *P* (respond “3” | *F*2, *B*), applying a log-linear correction for extreme proportions (Macmillan & Kaplan, 1985). From these, we derived the standard SDT measures of sensitivity (*d*^*′*^) and criterion (*c*) (Macmillan and Kaplan, 1985). Sensitivity and criterion were computed as *d*^*′*^ = *z*(*H*) − *z*(*FA*) and *c* = −0.5[*z*(*H*) + *z*(*FA*)].

*d*^*′*^ reflects the ability to discriminate two vs. three flashes, while *c* reflects the tendency to report “3”. *d*^*′*^ and *c* differences across beep levels (B0, B2, B3) were assessed using Friedman tests with Bonferroni-corrected Wilcoxon signed-rank post hoc comparisons. Within each beep condition, differences between the blind-spot and control locations were compared using Wilcoxon signed-rank tests (Bonferroni-corrected for multiple comparisons).

#### 2.5.5 Bayesian Causal Inference Model

We modeled participants’ multisensory responses using a Bayesian Causal Inference (BCI) framework, which estimates the probability that auditory and visual cues originate from a common source. Here, we will first describe the BCI model from which we will derive the forced-fusion, full-segregation and maximum likelihood estimation (MLE) models (Ernst and Banks, 2002) as alternative models. Details can be found in Körding et al. (2007).

The BCI model assumes that each sensory event *s*_*i*_ in the world causes a noisy sensation *x*_*i*_ of the event (Fig. 3A). Sensory noise is introduced by drawing *x*_*V*_ and *x*_*A*_ independently from Gaussian distributions centered on the true number of stimuli *s*_*V*_, *s*_*A*_ with standard deviations *σ*_*V*_, *σ*_*A*_ shown in Eq. 1.

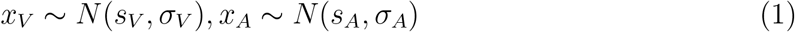

**Figure 3:**
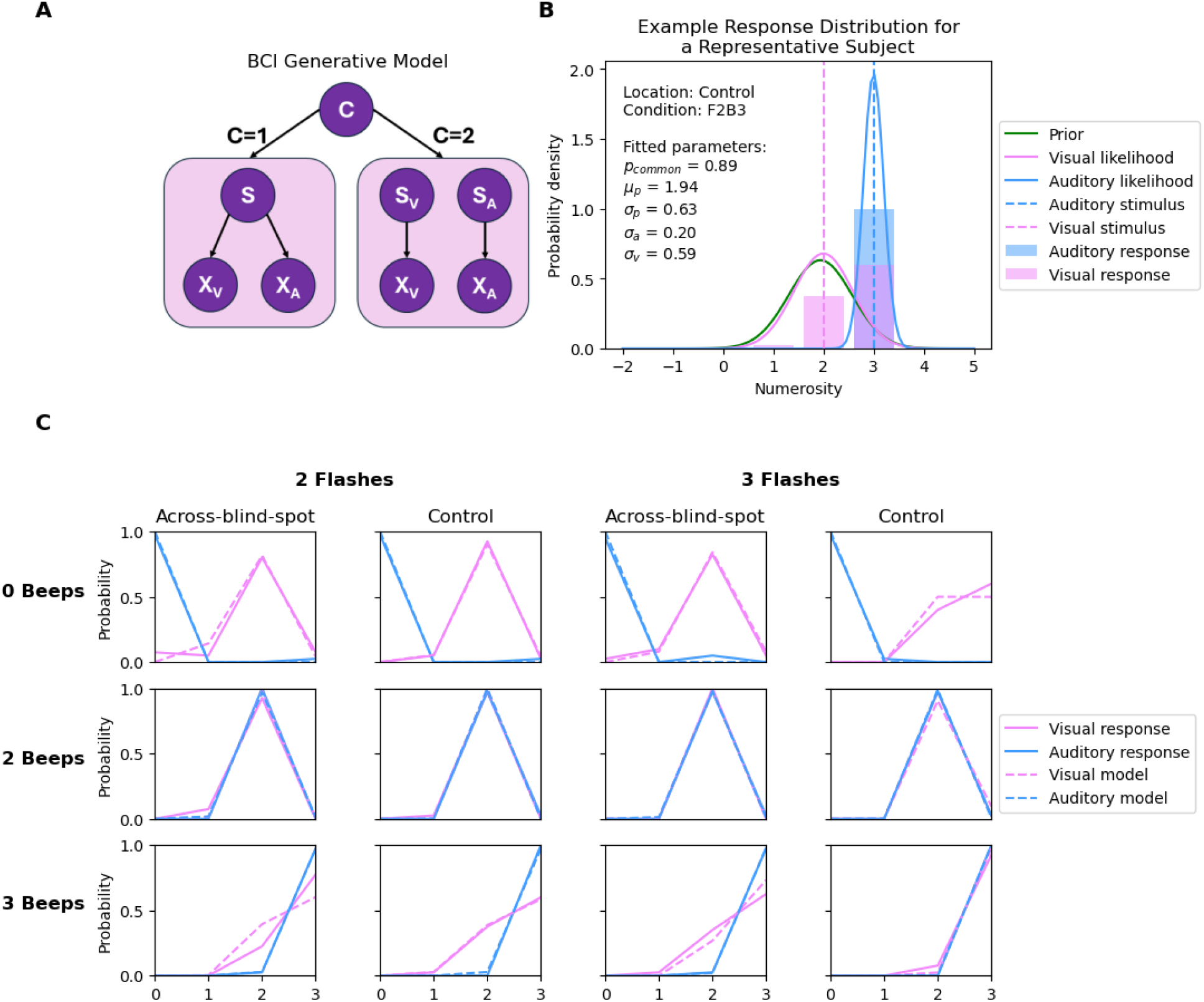
The generative model of the Bayesian Causal Inference model and model fitting results for a representative participant. **(A)** The BCI model where multisensory cues either share a common cause (C=1) or independent causes (C=2). If cues share a common cause, the final percept *s* is derived by cues from both modalities, i.e., *x*_*V*_ and *x*_*A*_, each perturbed by noises with width *σ*_*V*_ and *σ*_*A*_, respectively. If cues are independent, final percepts *s*_*V*_ and *S*_*A*_ are derived independently from *x*_*V*_ and *x*_*A*_. **(B)** The BCI model infers the causal structure by combining sensory likelihood and prior. Prior parameters include prior stimulus expectation (equation 4) and *p*_*common*_ (*a prior* expectation of a common cause). Sensory likelihood represents the sensory input (*x*_*V*_, *x*_*A*_) perturbed noise (*σ*_*V*_, *σ*_*A*_). Here, we have the fitted parameters of one representative participant and the predicted probability of auditory and visual responses. Since the participant has a high *p*_*common*_ and narrow auditory likelihood, the final percept *s* is heavily influenced by the auditory stimulus. textbf(C) Model fitting results for a representative participant. Solid pink and blue lines correspond to the visual and auditory response data, dashed pink and blue lines correspond to the model predictions for visual and auditory responses.

The model also assumes there is a prior bias for sensory information, modeled by a Gaussian distribution centered at *µ*_*P*_ with standard deviation *σ*_*P*_ . The prior distribution is given by Eq. 2:

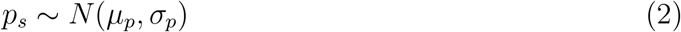

Common (*C* = 1) and independent (*C* = 2) causes are determined by sampling from a binomial distribution with the causal prior *p*(*C* = 1) = *p*_*common*_. The posterior probability of the underlying causal structure (unknown to the nervous system) can be inferred by combining the sensory evidence with causal prior according to Bayes rule:

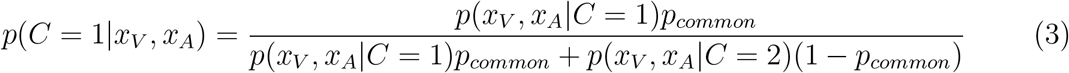

The causal prior quantifies observers’ tendency to assume a common cause before stimulus presentation. Post stimulus presentation, sensory evidence from both modalities informs the observers’ causal inference via the likelihood term.

If the sensory events can be attributed to a common cause, the estimate of *s* is a weighted average of both sensory signals and prior for *s*:

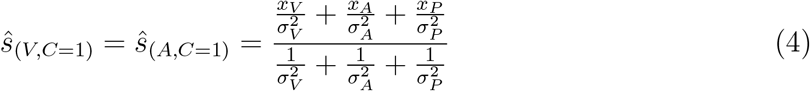

If sensory events are produced by independent causes, the estimate of *s* is a weighted average of unisensory signals and prior for *s*:

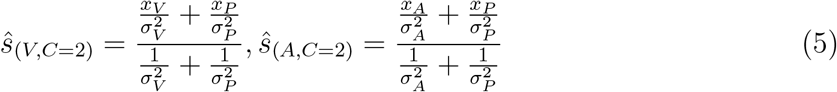

To obtain a final estimate of the number of visual or auditory stimuli, observers’ combine estimates under the two causal structures (*C* = 1, *C* = 2) using various strategies known as “model averaging”, “model selection”, and “probability matching”. Model averaging computes sensory estimates as the weighted sums of maximum likelihood estimates, with posterior probability as weights:

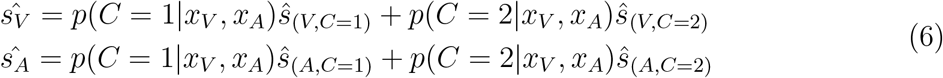

Model selection, as the name suggests, selects sensory estimates based on posterior probability — if *p*(*C* = 1|*x*_*V*_, *x*_*A*_) *>* 0.5, select *ŝ*_*C*=1_, select *ŝ*_*C*=2_ otherwise.

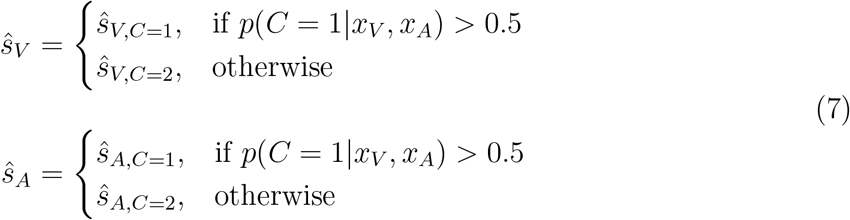

Probability matching is a stochastic decision making strategy where after calculating posterior probability, draw a random threshold *ξ* between [0, 1] on each trial, and select *ŝ* accordingly.

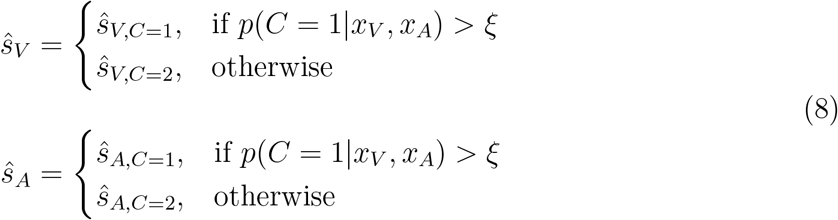

We evaluated model fits by comparing four models: (i) the full Bayesian Causal Inference (BCI) model, (ii) the forced-fusion model (i.e., formally, the BCI model with a fixed *p*_*common*_=1), (iii) the full-segregation model (i.e., formally, the BCI model with a fixed *p*_*common*_=0), (iv) the maximum-likelihood estimation (MLE) model, which assumes a common cause without prior information (i.e., formally, the BCI model with *p*_*common*_=1 and a flat prior, *σ*_*P*_ = 4000, *µ*_*P*_ = 1.5). An overview of these models is presented in Table 1. Because participants differed in their decision strategies (model averaging, model selection, or probability matching), and no single strategy dominated across participants (Supplementary Table S1), we used each participant’s best-fitting model for parameter estimation. The resulting parameter estimates were qualitatively similar across the three strategies.

**Table 1:** Overview of models and their free parameters.

| Model | Free Parameters | # |
| --- | --- | --- |
| Bayesian Causal Inference (BCI) | $p_{common}, \mu_P, \sigma_P, \sigma_V, \sigma_A$ | 5 |
| Forced-fusion | $\mu_P, \sigma_P, \sigma_V, \sigma_A$ | 4 |
| Full-segregation | $\mu_P, \sigma_P, \sigma_V, \sigma_A$ | 4 |
| Maximum-likelihood estimation (MLE) | $\sigma_V, \sigma_A$ | 2 |

Parameter estimation was performed using the BCI Toolbox (Zhu et al., 2024). Model comparison across strategies used the Bayesian Information Criterion (BIC) to identify the best-fitting model for each participant.

## 3 Results

### 3.1 Accuracy

As shown in Fig. 2A, for two-flash trials, accuracy significantly decreased when three beeps were presented, confirming that flash perception was significantly influenced by incongruent number of beeps presented (Control: *χ*^2^(2) = 29.34, *p <* .001; Across-blind-spot: *χ*^2^(2) = 17.62, *p <* .001). Similarly, for three-flash trials, accuracy decreased when only two beeps were presented, confirming that the auditory influence on flash perception (Control: *χ*^2^(2) = 31.68, *p <* .001; Across-blind-spot: *χ*^2^(2) = 32.65, *p <* .001).

Post hoc Wilcoxon signed-rank tests confirmed that bimodal incongruent conditions (F2B3, F3B2) yielded the lowest accuracy among all conditions (all Bonferroni-corrected *p <* .01). In contrast, bimodal congruent conditions (F2B2 and F3B3) produced higher accuracy than the corresponding unimodal conditions (F2B0 and F3B0; all Bonferroni-corrected *p <* .01). Notably, for three-flash trials, accuracy in the unimodal F3B0 condition was significantly lower than in the bimodal F3B3 condition across the blind spot, confirming that the middle flash was positioned within the blind spot and that F3B3 functionally resembled the F2B3 (Illusory AV Rabbit) configuration, with participants perceiving the ‘illusory’ (physically present) middle flash.

Beep-report accuracy remained near ceiling across all conditions and locations (mean accuracy *>* 0.9), confirming that participants reliably discriminated the number of auditory beeps. Because performance showed no meaningful variability across conditions, no further statistical analysis was conducted for beep responses.

### 3.2 Illusion Frequency

Illusion rates were comparable between control and across-blind-spot locations for both the Illusory Audiovisual Rabbit (F2B3) and the Invisible Audiovisual Rabbit (F3B2) conditions (all Bonferroni-corrected *p >* .05), indicating that the illusions occurred with similar strength in both spatial locations (Fig. 2B).

### 3.3 Perceived Flash Position

For each AV Rabbit illusion (F2B3, F3B2), we analyzed the distribution of reported flash positions (1–5) in illusion trials only (Fig. 2C). As expected for the AV Rabbit Illusion, reports were strongly non-uniform across conditions and locations, reflecting the characteristic shift toward the illusory intermediate position. Chi-square goodness-of-fit tests confirmed these patterned distributions (Illusory AV Rabbit (F2B3): Control *χ*^2^(4) = 815.27, *p <* .001; Across-blind-spot *χ*^2^(4) = 805.08, *p <* .001; Invisible AV Rabbit (F3B2): Control *χ*^2^(4) = 788.68, *p <* .001; Across-blind-spot *χ*^2^(4) = 779.16, *p <* .001).

The distributions of reported positions did not differ significantly between control and across-blind-spot locations (Illusory AV Rabbit (F2B3): *χ*^2^(4) = 8.60, *p* = .072; Invisible AV Rabbit (F3B2): *χ*^2^(4) = 3.46, *p* = 0.48), indicating that both illusions exhibited comparable spatial patterns across locations.

Consistent with the numbered location reports, click-based reports of illusory and last-flash locations collected in the *Additional Flash Localization Task* were centered near the corresponding reference locations for both control and across-blind-spot conditions (Fig. 2E). Two-dimensional kernel density maps revealed overlapping clusters of perceived illusory and last-flash positions when aligned to the respective reference points (blind-spot/control center or true last-flash location), suggesting comparable localization precision across locations. The mean distance between perceived illusory flashes and the reference center was 1.38^°^(*SD* = 1.06^°^), whereas the mean distance between perceived last flashes and their true physical locations was 1.79^°^(*SD* = 1.40^°^), indicating modest variability in spatial accuracy across trials.

### 3.4 Signal Detection Theory

A Friedman test revealed a significant effect of number of beeps on sensitivity (*d*^*′*^) at control locations (*χ*^2^ = 27.1, *p <* .001), but not across the blind spot (*χ*^2^ = 2.96, *p* = 0.23) (Fig. 2D. Post hoc Wilcoxon signed-rank tests (Bonferroni-corrected) showed that at control locations, sensitivity was significantly lower for both the two-beep (B2) and three-beep (B3) conditions compared with the visual-only (B0) condition (all *p <* .001), while no difference was observed between the B2 and B3 conditions (*p* = 0.81). At across-blind-spot locations, *d*^*′*^ did not differ significantly across beep contexts (all *p >* .1).

Beep context strongly modulated the response criterion (*c*) but not sensitivity (*d*^*′*^). Friedman tests showed a significant main effect of beep number on criterion in both control and across-blind-spot locations (*χ*^2^ = 40, *p <* .001 for each). Post hoc Wilcoxon signed-rank tests (Bonferroni-corrected) confirmed that criterion values were highest for the two-beep condition (B2), lowest for the three-beep condition (B3), and intermediate for the visual-only condition (B0; all *p <* .001). This pattern indicates a systematic response bias, with two beeps biasing reports toward two flashes and three beeps biasing reports toward three flashes.

We compared sensitivity (*d*^*′*^) and criterion (*c*) between control and across-blind-spot locations within each beep condition (B0, B2, B3) using Wilcoxon signed-rank tests with Bonferroni correction. Sensitivity was higher at control than across-blind-spot locations for the unimodal baseline (B0; *W* = 0, *p <* .001) and Invisibe AV Rabbit context (B2; *W* = 23, *p* = .011). The Illusory AV Rabbit context (B3) showed a similar trend that did not survive correction (*W* = 48, *p* = 0.098). In contrast, response criterion was consistently more conservative across the blind spot for all beep contexts (B0: *W* = 6, *p <* .001; B2: *W* = 34, *p* = 0.042; B3: *W* = 28, *p* = 0.008), indicating a greater tendency to report “2” across the blind spot.

Although Wilcoxon tests indicated that response criterion (*c*) was slightly higher across the blind spot than at control locations in each beep condition (all *p <* .05), the mean absolute differences were small in magnitude (Δ = 0.18 − 0.37) and accompanied by substantial overlap of confidence intervals (Fig. 2D). These results suggest a very small criterion offset across the blind spot, with no evidence of a qualitatively different decision strategy.

Together, these results show that the blind spot reduces visual discriminability and shifts decision bias away from “3,” whereas the number of beeps primarily modulates response bias (see Friedman analyses), consistent with the AV Rabbit effects.

### 3.5 BCI Modeling Results

#### 3.5.1 Model Comparison

We compared model performance across the Bayesian Causal Inference (BCI; see Fig. 3A), forced-fusion, full-segregation, and Maximum Likelihood Estimation (MLE) models using the Bayesian Information Criterion (BIC). For each participant and stimulus location, the model with the lowest BIC score was considered to provide the best trade-off between goodness of fit and model complexity.

Across participants, the BCI model consistently yielded lower BIC scores than forced-fusion, full-segregation, and MLE models (Across-blind-spot: 160.28 *±* 12.4; Control: 131.47 *±* 10.9), indicating superior explanatory power. The corresponding forced-fusion (Across-blind-spot: 210.56 *±* 20.49; Control: 159.67 *±* 15.37), full-segregation (Across-blind-spot: 225.85 *±* 21.55; Control: 191.37 *±* 13.70), and MLE (Across-blind-spot: 245.33 *±* 23.71; Control: 183.02 *±* 19.02) models fit the data substantially worse.

#### 3.5.2 BCI Model Estimated Parameters

We fitted a Bayesian Causal Inference model with five free parameters for each participant, stimulus location (across-blind-spot and control), and decision strategy (model averaging, model selection, and probability matching). For each participant and location, the best-fitting strategy was identified by comparing Bayesian Information Criterion (BIC) values across the three models. Table 2 summarizes the estimated parameters from these best-fitting models (see Fig. 3B for the visualization of fitted parameters for a representative participant). Because participants differed in their preferred strategies and no single strategy consistently provided the best fit (Supplementary Table 1), parameter estimates were taken from each participant’s individually best-fitting model. Overall, the estimated parameters were qualitatively similar across strategies.

**Table 2:** Parameter estimates (mean *±* SE) across participants and stimuli locations.

| Location | Parameters |  |  |  |  |
| --- | --- | --- | --- | --- | --- |
| | $p_{common}$ | $\sigma_P$ | $\mu_P$ | $\sigma_V$ | $\sigma_A$ |
| Across-blind-spot | $0.50 \pm 0.06$ | $1.19 \pm 0.13$ | $1.90 \pm 0.13$ | $2.35 \pm 0.12$ | $0.21 \pm 0.02$ |
| Control | $0.58 \pm 0.08$ | $0.85 \pm 0.07$ | $2.32 \pm 0.16$ | $1.34 \pm 0.15$ | $0.20 \pm 0.01$ |

Goodness of fit was quantified using the coefficient of determination (*R*^2^) between the model-estimated numerical responses and the participant-reported numerical responses (i.e., perceived number of beeps and flashes). The best BCI models demonstrated excellent per-formance, achieving a mean *R*^2^ of 0.99 across all 20 participants (see Fig. 3C for model fitting results for a representative participant).

Wilcoxon signed-rank tests revealed significant location-dependent differences in prior and visual parameters (*σ*_*P*_ : *W* = 16, *p* = 0.0016; *µ*_*P*_ : *W* = 18, *p* = 0.0024; *σ*_*V*_ : *W* = 9, *p <* .001). No significant differences were found for the causal prior *p*_*common*_ (*W* = 49, *p* = 0.18) and auditory parameter *σ*_*A*_ (*W* = 81, *p* = 1.00).

Together, these findings indicated greater visual uncertainty and wider (looser) prior stimulus expectations across the blind spot compared to control locations, whereas auditory reliability and causal prior remained stable.

## 4 Discussion

In this work, we asked whether multisensory filling-in extends into the physiological blind spot, and if so, how this process is computationally achieved. We asked three questions: (i) Does audiovisual integration produce the Audiovisual Rabbit Illusion (Stiles et al., 2018) across the blind spot? (ii) If so, is the illusion enhanced because the brain has a stronger tendency to fill in missing information where vision is absent? (iii) Even if behavior appears similar across regions, are the underlying computational mechanisms the same or fundamentally altered in the blind spot? Our findings address these questions in turn.

### Does multisensory filling-in occur across the blind spot?

Yes. Participants experienced robust Audiovisual Rabbit Illusions in both the classic (F2B3) and invisible (F3B2) variants across the blind spot (Fig. 2A). This demonstrates that audio-visual binding does not require direct visual input from the interpolated region. Instead, the percept appears to depend on internal generative models that enforce spatial and temporal continuity. Prior work on filling-in has largely emphasized unimodal mechanisms (Komatsu, 2006; Devinck and Knoblauch, 2019), or compensatory enhancements in long-term blindness (Kupers and Ptito, 2014; Braun, 2016). Our results extend these insights to multisensory perception in sighted observers: the brain treats the blind spot as part of a continuous visual scene and incorporates auditory structure to construct a percept that is not directly supported by retinal input.

Localization data reinforce this conclusion. Both number-based and click-based reports showed that perceived flashes—illusory or real—clustered near their expected spatial positions (Fig. 2C, E). Mean localization errors remained within a few degrees, indicating that spatial interpolation at the blind spot preserves the coordinate structure necessary for multisensory alignment.

### Is the illusion enhanced across the blind spot due to stronger filling-in?

No. Contrary to the possibility that missing visual information might amplify reliance on auditory cues, illusion strength was comparable across blind-spot and control regions (Fig. 2B). This suggests that the brain does not default to a stronger common-cause assumption when visual evidence is absent. In other words, the blind spot does not behave like a “perceptual vacuum” in which crossmodal signals automatically dominate.

This finding constrains interpretations of sensory compensation. Although compensatory enhancements have been documented in long-term or congenital visual deprivation (Ptito et al., 2012), our results show that compensation at the blind spot is not expressed as increased multisensory fusion. Instead, the blind spot is integrated using the same overall inference structure as visible regions.

### Are the underlying computational mechanisms preserved or altered?

Here we find a dissociation: the causal inference architecture is preserved, but the sensory and prior uncertainties differ.

Bayesian Causal Inference (BCI; Fig. 3A) provided the best quantitative account of behavior at both locations, outperforming forced-fusion, full-segregation, and MLE models. This replicates prior demonstrations that observers dynamically arbitrate between common- and separate-cause interpretations (Körding et al., 2007; Shams and Beierholm, 2022), and shows that this flexible inference is maintained even where bottom-up vision is absent.

At the same time, key fitted parameters diverged across locations (Table 2): (i) Causal prior (*p*_*common*_) was comparable across blind-spot and control regions. Participants were therefore not more likely to infer a common cause simply because the visual signal was unreliable. (ii) Auditory likelihood (*σ*_*A*_) was also unchanged, indicating that participants relied on auditory cues with similar precision in both across blind spot and control regions (iii) Visual likelihood (*σ*_*V*_) was broader across the blind spot, consistent with greater visual uncertainty when there is no bottom-up retinal input from the blind spot. (iv) Numerosity prior (*σ*_*P*_) was flatter in the blind spot, indicating weaker expectations about intermediate event counts. (v) Prior mean (*µ*_*P*_) was shifted lower in across-blind-spot trials, suggesting reduced expectation for number of visual events.

Overall, this data supports the notion that the internal causal structure remains stable across regions, in agreement with Odegaard and Shams (2016) ‘s findings, where binding tendency is stable within a task and across time, but is not generalized across tasks. The flatter numerosity prior across blind-spot trials also aligns with evidence that the Audiovisual Rabbit depends on predictive and postdictive processes that integrate information over several hundred milliseconds (Stiles et al., 2018). In the absence of bottom-up vision, the system may distribute its expectations more broadly and rely more heavily on temporally extended context, leading to altered prior precision without changes to causal architecture. This separation between architecture (preserved) and parameterization (adjusted) is a central theoretical contribution of the present work. It shows that multisensory inference tolerates missing modality-specific evidence by adapting uncertainty rather than altering causal expectations.

### Overall significance

Together, these results characterize how the brain fills in the blind spot in multisensory contexts. The perceptual system maintains continuity not by increasing binding tendencies, but by applying a stable causal inference framework and recalibrating sensory and prior uncertainties to reflect the absence of local signals. Multisensory perception persists through probabilistic reconstruction rather than direct sensory data, revealing how the brain sustains coherent perception despite structural gaps in input.

## Acknowledgements

We are grateful for support from the National Institutes of Health, National Eye Institute (R01EY031761); the National Institutes of Health, the BRAIN Initiative (5K99EY031987-02). A.C. thanks the Croucher Foundation for a scholarship.

## Author Contributions

Ailene Chan: Conceptualization; Methodology; Software; Validation; Formal Analysis; Investigation; Visualization; Project Administration; Writing – Original Draft; Writing – Review & Editing

Noelle Stiles: Methodology; Funding Acquisition; Supervision; Writing – Review & Editing Carmel Levitan: Formal Analysis; Funding Acquisition; Supervision; Writing – Review & Editing

Armand Tanguay, Jr: Methodology; Funding Acquisition; Supervision; Writing – Review & Editing

Daw-An Wu: Methodology; Writing – Review & Editing

Shinsuke Shimojo: Methodology; Funding Acquisition; Supervision; Writing – Review & Editing

## Data and code availability statement

The figures in this article, as well as the data and plotting scripts necessary to reproduce them, are available openly under the CC-BY license (Chan, 2025). Additionally, custom scripts developed for the experiments are included in this release.

