## Supplementary Information for "Bayesian Causal Inference Accounts for Multisensory Filling-In at the Blind Spot"

**Figure S1. Blind Spot Measurements (N = 20)**

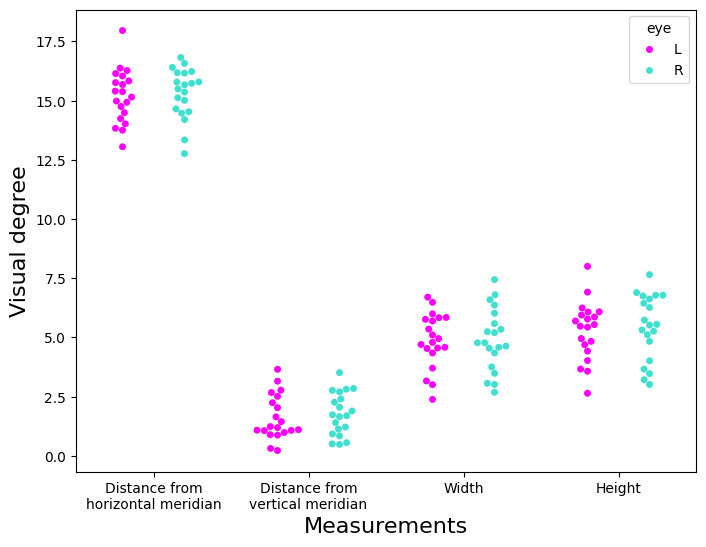

The distribution of distances from the vertical and horizontal meridians, as well as the width and height of the blind spots for each eye (left eye: magenta, right eye: turquoise). Here, the vertical and horizontal meridians refer to the midlines of the monitor along the x- and y-axes. Blind-spot locations and sizes were consistent across individuals. The mean blind-spot area was 21.6 deg^2^ (SD = 8.99). Blind-spot centers were located approximately 15.2° from the fovea (SD = 0.976) and 1.69° below the horizontal meridian (SD = 0.780).  **Supplementary Figure S2. Eye-tracking measurements during fixation, stimulus presentation, and response phases (N = 20)**

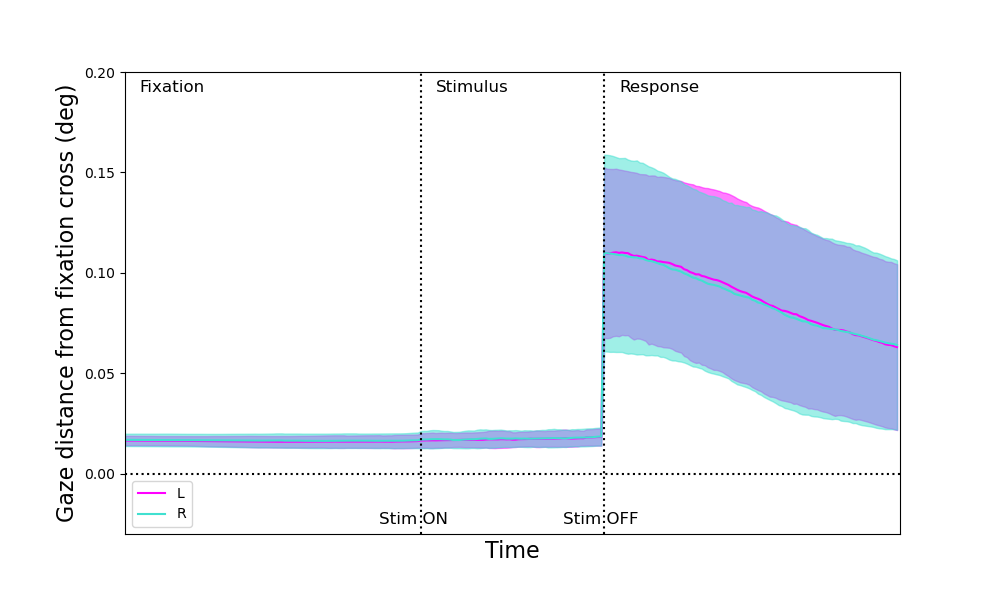

The y-axis represents the gaze-point distance from the fixation cross (center) in visual degrees, and the x-axis represents time. Solid lines indicate the mean gaze distance across trials, and shaded regions indicate standard deviations. Color code: left eye – magenta; right eye – turquoise. Participants maintained excellent fixation throughout the fixation and stimulus presentation phases. Mean gaze-point distance from the center was 0.016° (SD=0.0028) for the left eye and 0.017° (SD=0.0034) for the right eye during fixation, and 0.017° (SD=0.0039) and 0.018 (SD=0.0043) during stimulus presentation, respectively. Although gaze deviations increased slightly during the response phase (e.g., when reading the text prompt or locating response keys), fixation remained stable during stimulus presentation, confirming that flashes were presented accurately at the intended retinal locations (borders and center of the blind spots). Individual eye-tracking data for all participants are shown in Supplementary Figure S3. **Supplementary Figure S3. Individual eye tracking measurements during fixation, stimulus presentation, and response phases**

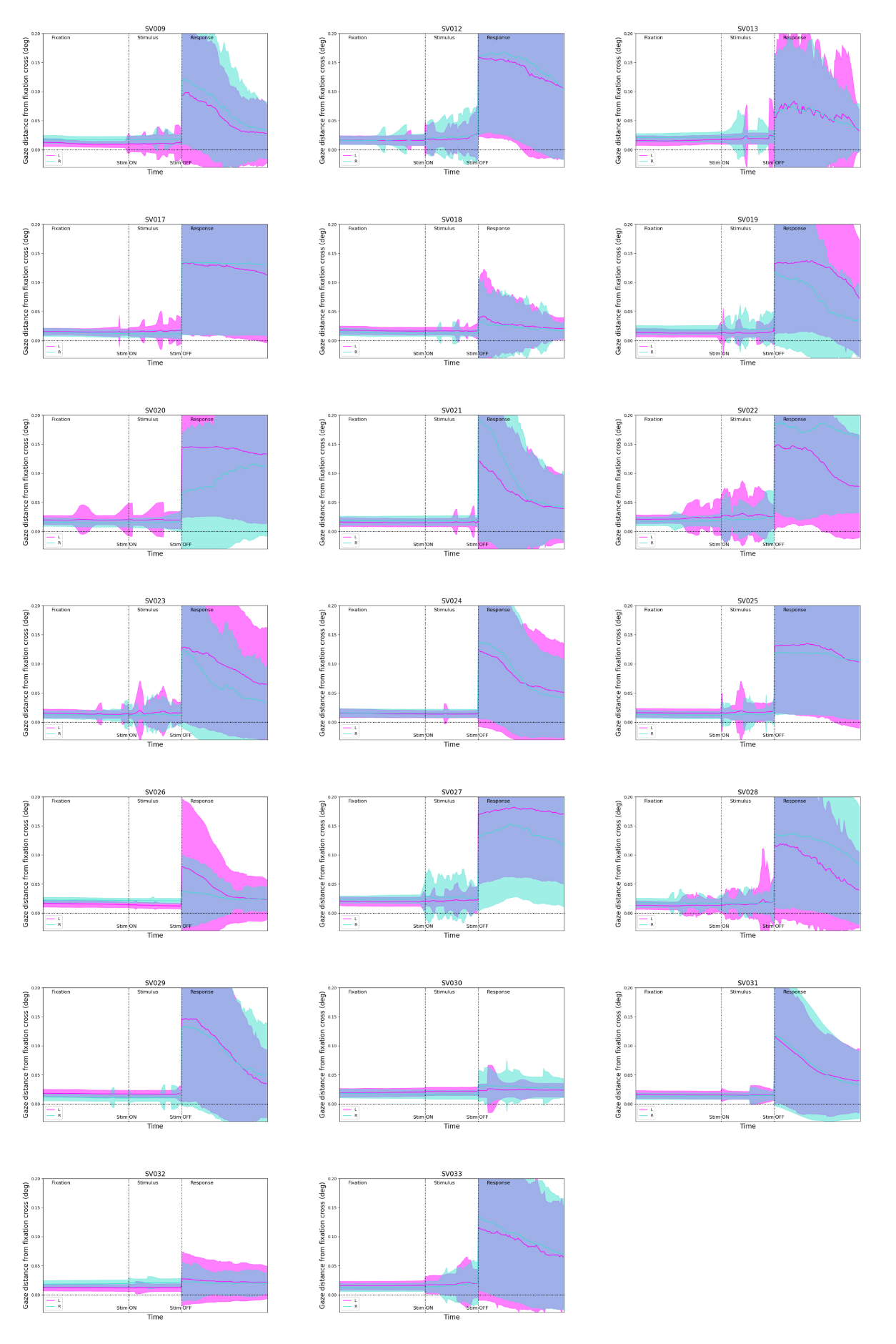

**Supplementary Table S1. Comparison between three decision strategies of the BCI model (model averaging, model selection, and probability matching).**

| Strategy | Location | *p*_common_ | *µ*_P_ | *σ*_p_ | *σ*_v_ | *σ*_A_ | *R*^2^ | BIC | % win |
| --- | --- | --- | --- | --- | --- | --- | --- | --- | --- |
| Model Averaging | Across-blind-spot | 0.55 ± 0.07 | 1.67 ± 0.17 | 1.12 ± 0.15 | 1.65 ± 0.19 | 0.25 ± 0.03 | 0.96 ± 0.02 | 184.43 ± 17.41 | 45.0 |
|  | Control | 0.59 ± 0.08 | 2.28 ± 0.17 | 0.87 ± 0.08 | 1.12 ± 0.14 | 0.33 ± 0.13 | 0.95 ± 0.03 | 146.64 ± 15.38 | 45.0 |
| Probability Matching | Across-blind-spot | 0.49 ± 0.08 | 1.76 ± 0.16 | 1.09 ± 0.12 | 1.79 ± 0.20 | 0.20 ± 0.02 | 0.97 ± 0.01 | 167.94 ± 12.19 | 20.0 |
|  | Control | 0.59 ± 0.08 | 2.26 ± 0.18 | 0.82 ± 0.07 | 1.12 ± 0.15 | 0.19 ± 0.01 | 0.98 ± 0.00 | 135.07 ± 10.33 | 30.0 |
| Model Selection | Across-blind-spot | 0.49 ± 0.05 | 1.91 ± 0.14 | 1.05 ± 0.14 | 1.92 ± 0.16 | 0.23 ± 0.02 | 0.97 ± 0.01 | 166.67 ± 13.16 | 35.0 |
|  | Control | 0.60 ± 0.06 | 2.33 ± 0.16 | 0.74 ± 0.08 | 1.05 ± 0.11 | 0.20 ± 0.01 | 0.98 ± 0.00 | 133.99 ± 11.09 | 25.0 |

Note: ***p*_common_**, causal prior; ***µ*_P_**, mean of the numeric prior; ***σ*_p_**, standard deviation of the numeric prior; ***σ*_v_**, standard deviation of the visual likelihood; ***σ*_A_**, standard deviation of the auditory likelihood; ***R*^2^**, coefficient of determination; **% win**, percentage of participants in which a model won the within-participant model comparison based on BIC.
